# Disparate effects of antibiotic-induced microbiome change and enhanced fitness in *Daphnia magna*

**DOI:** 10.1101/586669

**Authors:** Asa Motiei, Björn Brindefalk, Martin Ogonowski, Rehab El-Shehawy, Paulina Pastuszek, Karin Ek, Birgitta Liewenborg, Klas Udekwu, Elena Gorokhova

## Abstract

It is a common view that an organism’s microbiota has a profound influence on host fitness; however, supporting evidence is lacking in many organisms. We manipulated the gut microbiome of *Daphnia magna* by chronic exposure to different concentrations of the antibiotic Ciprofloxacin (0.01 – 1 mg L^−1^), and evaluated whether this affected the animals’ fitness and antioxidant capacity. In line with our expectations, antibiotic exposure altered the microbiome in a concentration-dependent manner. However, contrary to these expectations, the reduced diversity of gut bacteria was not associated with any fitness detriment. Moreover, the growth-related parameters correlated negatively with diversity indices; and, in the daphnids exposed to the lowest ciprofloxacin concentrations, the antioxidant capacity, growth, and fecundity were even higher than in control animals. These findings suggest that ciprofloxacin exerts direct stimulatory effects on growth and reproduction in *Daphnia*, while microbiome-mediated effects are of lesser importance. Thus, although microbiome profiling of Daphnia may be a sensitive tool to identify early effects of antibiotic exposure, disentangling direct and microbiome-mediated effects on host fitness is not straightforward.

## Introduction

In multicellular organisms, the microbiome contributes to critical aspects of host development and physiology (1). In ecological, evolutionary and ecotoxicological research, there is growing recognition that environmental stresses imposed upon the microbiome may drive physiological responses, life-history penalties and adaptation capacity of their hosts (2), (3) 4). Consequently, coping with various environmental insults would involve both the host and its microbiome responses.

The gut microbiota participates directly in food digestion and nutrient assimilation, which affects the host’s energy acquisition and growth (5). In addition to this, the host immune system is influenced by the gut microbes via a number of different mechanisms, e.g., competition with pathogens as well as suppression and modification of virulence factors via metabolite production (6). Symbiotic bacteria are also capable of enhancing the host innate immune system by, for example, up-regulation of mucosal immunity, induction of antimicrobial peptides and antibodies (7, 8). Considering the biological effects triggered by the host-microbiome interactions, a disruption of mutualistic bacterial communities may result in increased susceptibility to pathogens and infections, while simultaneously affecting the growth and development of the host via compromised nutrition. In various gnotobiotic animal models, poor survival, growth and fecundity are commonly observed, reflecting a physiological impairment due to some dysbiotic state of microflora (3, 9).

If growth penalties are to be expected in animals with perturbed microbiota, then it should be possible to manipulate animal fitness by targeting its resident bacteria with antibacterial substances. In line with this, retarded development has been observed in the copepod *Nitocra spinipes* upon antibiotic exposure, and linked to structural changes in its microbiota (10). It was suggested that aberrant digestion was behind these changes as has also been observed in *Daphnia magna* following a short-term antibiotics exposure (9,11). Moreover, an altered microbiota composition was reported in *Daphnia* following a long-term exposure to the antibiotic oxytetracycline, concurrent with reduced host growth (12). While perturbed microbiota can manifest itself directly as decreased nutrient uptake, another outcome can be effects on host antioxidant production, with concomitant effects on immunity and growth (13). However, short antibiotic exposure may not necessarily result in any significant growth penalties in the long run. The outcome of any chronic exposure to antibiotics would largely depend on the resilience of the bacterial communities, and their capacity to recover and re-establish any functional interaction(s) relevant to the host (16,17,18,19,20).

To study the relationships between microbiome composition and host performance, a common set of model species and methods to manipulate their microbiomes is needed. In ecology, evolution and ecotoxicology, *Daphnia* species are used routinely as model organisms because of their well-known physiology, rapid reproduction, and sensitivity to environmental factors (19). The microbiome of the laboratory-reared *Daphnia magna* has been recently presented in several studies using different approaches, from cloning to shotgun sequencing (22, 23). Regardless of the sequencing platform, origin of specimens, and culture conditions, the core microbiome appears relatively stable, mainly comprised of *Betaproteobacteria, Gammaproteobacteria* and facultative anaerobic *Bacteroidetes* species. At the genus level, *Limnohabitans* has been reported as one of the most stable and dominant members in *Daphnia* gut, and variations in its abundance have been tied to the animal fecundity (22). Although some studies have addressed the dependence of *Daphnia* on its microbiota (9) and some short-term effects on fitness following exposure to antibiotics have been observed in *Daphnia magna* (25, 13), the relationship between microflora perturbation and host fitness is still unclear, as is the involvement and modulating role of antioxidants in these relationships.

In this study, the relationship between antibiotic-mediated gut microbiome modulation and host fitness were addressed experimentally using a model cladoceran *Daphnia magna*. We monitored changes in the gut microbiome, host longevity, growth, and reproduction, as well as antioxidant levels in the exposed animals following ciprofloxacin exposure. We hypothesized that the diversity and abundance of the gut-associated microflora would decrease with increasing concentration of antibiotics. Furthermore, we expected longer exposure time and higher antibiotic concentrations to have negative effects on somatic growth, reproductive output, and antioxidant capacity. These reductions we expected would be due to reduced bacterial diversity in particular, and to some extent, an altered community composition. These hypotheses were tested by combining (1) long-term (21 d) exposure experiments with life-table analysis, (2) microbiome profiling using the next generation sequencing of 16S rRNA gene and taxonomic assignment, and (3) measurements of daphnid total antioxidant capacity, growth, and fecundity.

## Material and methods

### Test species and culture conditions

The cladoceran *Daphnia magna*, originating from a single clone (Environmental pollution test strain *Clone 5*, Federal Environment Agency, Berlin, Germany), was used in this experiment. The animals were cultured in groups of 20 individuals in 3-L beakers with M7 medium (OECD standard 202 and 211), and fed a mixture of the green algae *Pseudokirchneriella subcapitata* and *Scenedesmus subspicatus* three times a week; the algae were grown axenically.

### Ciprofloxacin stock solutions

Ciprofloxacin hydrochloride (CAS: 86393-32-0; Sigma) the antibiotic utilized in this study and is a broad spectrum fluoroquinolone, active against both Gram-positive, G+, and Gram-negative, G-, bacteria. Its mode of action is the inhibition of the gyrase and / or topoisomerase enzyme of microbes which determines the supercoiling state of DNA, and critical to bacterial replication, repair, transcription and recombination (24). Selection of this drug was due to its rapid absorption, long half-life in the test system, and the absence of acute toxicity in *D. magna* within the range of the concentrations tested (25). A singular stock solution of ciprofloxacin (1 mg/ml) was prepared in M7 medium and stored at -20°C during the course of the experiment.

### Experimental design

We employed three drug concentrations (0.01, 0.1 and 1 mg/L) and a control treatment (M7 medium). For each treatment, 25 neonates (< 24 h) of *D. magna* were placed individually in 40 mL of M7 medium, with or without ciprofloxacin; the medium was changed every second day. The test design followed a standard procedure for the reproduction test with *Daphnia* (OECD standard 211). The animals were fed daily with a suspension of green algae *Pseudokirchneriella subcapitata* (0.2 mg C d^−1^; axenic culture) and incubated at 22°C with 16^L^: 8^D^ photoperiod. Under these conditions, the animals matured and started to reproduce 8-9 d after the start of the experiment. All jars were inspected daily and mortality recorded. Upon release of neonates, counts were made, offspring discarded, and brood size recorded for each female and within each brood. In conjunction with brood release, four randomly selected individuals from each treatment were placed in antibiotic-free medium. In this manner, we collected females after their 1^st^, 2^nd^, 3^rd^, and 4^th^ clutch, with the last individuals sacrificed on day 21, when the experiment was terminated. When sampling, the images of females were acquired by scanning live animals on a glass surface in a drop of water (CanoScan 8800F 13.0), and their body length (BL, mm) was measured using ImageJ software (26). For each individual, the gut was dissected using a sterile needle and a pair of forceps, washed with nuclease-free water, transferred individually to Eppendorf tubes and stored at -80°C until DNA extraction. The degutted body was transferred to a fresh Eppendorf tube, stored at -80°C and tested for antioxidant levels based on Oxygen Radical Absorbance Capacity (ORAC) and protein content.

### DNA Extraction

DNA was extracted from the gut samples using 10% Chelex (24) and purified with AMPure^R^ XP beads (Beckman Coulter, Brea, CA, USA) following the manufacturer’s instructions. Initial DNA concentrations following purification were evaluated using Quant-iT PicoGreen dsDNA Assay kit (ThermoFisher, USA) following the instructions from L(27)). Absorbance was measured at 530 nm, using a Tecan Ultra 384 SpectroFluorometer (PerkinElmer, USA).

### 16S rRNA gene amplification and sequencing library preparation

Bacterial diversity of the samples was analyzed by sequencing of amplicons generated from the V3-V4 region of the 16S rRNA gene using the MiSeq Illumina platform. Two-stage PCR amplification was performed using forward primer 341F: (CCTACGGGNGGCWGCAG) and reverse primer 805R: (GGACTACHVGGGTWTCTAAT). The first PCR was carried out in 25-µl PCR reactions and comprised 0.02 U µl^−1^ Phusion polymerase (ThermoFisher, USA), 0.2 mM dNTP, 1 mM MgCl_2_, 1 × Phusion reaction buffer, 0.5 µM of each primer as well as 5 ng of DNA template). The amplification protocol consisted of an initial denaturation at 98 °C for 30 seconds followed by 35 cycles of 10 sec at 98 °C, 30 sec at 55 °C and 72 °C, and, a final extension step (72 °C for 10 min). PCR products were purified using Agencourt AMPure XP beads (Beckman Coulter, Brea, CA, USA). Following this, amplicon PCR was performed on 5 µl of equimolar amounts of PCR product using Nextera XT primers (Index 1 (N7XX) and Index 2(S5xx)), targeting the same region of the 16S rRNA genes (8 cycles of 30 sec at 95 °C, 30 sec at 55 °C and 35 sec at 72 °C). The products were purified with Amplicons AMPure XP Beads (Beckman Coulter) according to the manufacturer’s protocol and concentrations estimated using Quant-iT PicoGreen dsDNA Assay kit (ThermoFisher, USA). Individually barcoded samples were mixed in equimolar amounts, and DNA sequencing adaptor indexes ligated using the TruSeq DNA PCR-free LT Library Preparation Kit (Illumina). Quality control was performed on an Agilent 2100 BioAnalyser using high sensitivity DNA chip. PhiX DNA (10%) was added to the denatured pools, and sequencing was performed on an Illumina MiSeq using the MiSeq V3 reagent kit (600-cycles) on the Illumina MiSeq platform. De-multiplexing and removal of indexes and primers were done with the Illumina software v. 2.6.2.1 on the instrument according to the standard Illumina protocol.

### Processing of sequencing data

Following initial upstream de-multiplexing and index removal, sequences were analysed using the *DADA2* v. 1.6 module (28) as implemented in the R statistical software v. 3.4.2 (29). The pipeline consisted of quality-filtering, trimming of bad quality (< Q30) stretches, error estimation and de-replication of reads, merging of forward and reverse reads and finally, removal of chimeric sequences. All remaining sequences were assigned taxonomy on the genus level using the Silva Ribosomal RNA database version v.128. Subsequent statistical analyses and visualization were done with the *Phyloseq* R-module v.1.22.3 (30) unless otherwise stated.

### Analysis of Oxygen Radical Absorbance Capacity and protein content

As a proxy for antioxidant capacity, we assayed oxygen radical absorbance capacity (ORAC) according to (31) with minor modifications and normalized values to protein content. This biomarker represents the water-soluble fraction of antioxidants and has been applied in daphnids (32). Samples for ORAC and protein measurements were homogenized in 100 μL of PPB buffer (75 mM, pH 7.4). Fluorescein was applied as a fluorescent probe (106 nM) and 2, 2-azobis (2-amidinopropane) dihydrochloride (AAPH) (152.66 mM) as a source of peroxyl radicals. Trolox (218 μM, Sigma–Aldrich) was used as the standard. The assay was conducted in 96-well microplates while 20 μL of homogenate sample was added to each well and mixed with 30 μL of AAPH and 150 μL of fluorescein. Fluorescence was measured at 485nm/520nm. Protein content of the supernatant was determined by the bicinchoninic acid method using a Pierce BCA Protein Assay kit 23227 (Thermo Scientific) according to the microplate procedure with some modifications. In each well, 25 µl of blank, standard or samples was added to 200 µl of working solution. Absorbance was measured at 540 nm using a FluoStar Optima plate reader (BMG Lab Technologies, Germany). Antioxidant capacity was expressed as mg trolox eq. mg protein^−1^.

## Data analysis and statistics

### Life-history traits

Survival probability was calculated using Kaplan-Meier analysis, which estimates the probability of an event (i.e., death) occurring in a given period (33). The logrank test was used to evaluate differences in the survivorship between the treatments using package *survival* in R (34).

The empirical von Bertalanffy growth model was applied to determine growth parameters using length-at-age data fitted to the equation:

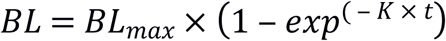

where *BL* is the total length at time *t* (days); *BL*_*max*_ is the length reached at an infinite time, defined as the maximum potential length attained under the prevailing conditions; and *K* is the growth rate. Statistical differences in *BL*_*max*_ and *K* between each treatment and control were determined by non-overlapping 95% confidence intervals.

To analyze the effects of exposure time and ciprofloxacin concentration on the daphnid fecundity, we used generalized linear models (GLM) with Poisson distribution and identity link function. Residuals were checked visually, and nonsignificant interaction terms were dropped from the analysis. A post hoc Tukey HSD test was used to compare the brood size among the treatments for each clutch.

The daphnid population growth rate (PGR) was estimated according to Euler-Lotka’s equation using (R Core Team, 2018) (Appendix S10):

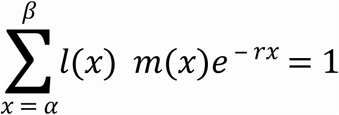

where *l(x)* is the fraction of individuals surviving to age *x* and *m(x)* is the birth rate per capita for the mothers of age *x*. Bootstrapping (999 permutations) was used to estimate 95% confidence limits of the PGR values in each treatment, and statistical differences in *r* between each treatment and control were determined by non-overlapping 95% confidence intervals.

### Analysis of microbiota communities

#### Diversity indices analysis

To assess the alpha diversity of the bacterial communities, we calculated commonly used indices of diversity and evenness (ACE, Chao1 and Fisher’s alpha). Effects of time and concentration on the diversity indices were tested by GLM with normal error structure and log-link. Quantile plots were used to evaluate the distribution of the residuals and deviance was used to access goodness of the model. Interaction (*time* × *concentration*) was first included in every model but omitted if found not significant. The Principal coordinates analysis (PCoA) with Bray-Curtis dissimilarity index was used to visualize differences in community composition among the treatments (35). Differences in the community structure at the family level were tested by permutational multivariate analysis of variance (Permanova) using variance stabilized Bray-Curtis dissimilarity. Multivariate homogeneity of treatment dispersion was assessed using the beta-disperser in the *vegan* package (36).

#### Connecting the microbiome to host fitness

The R-package *edgeR* (37) was used to identify differentially abundant bacterial taxa (false discovery rate-corrected *P*-values, α = 0.05, FDR=1%) that were associated with high or low growth rate (somatic and reproductive) of the daphnids. As a measure for somatic and reproductive growth, we used BL and fecundity, respectively. For each trait, we created two classes, *high* (above the group mean, coded as 1) and *low* (below the group mean, coded as 0) using zeta scores for individual BL and fecundity measurements. Zeta scores (zero mean, unit variance) were calculated based on clutch-specific mean values (all treatments included) and corresponding standard deviations to account for the changes in BL and fecundity with the daphnid age.

## Results

### Survival and individual growth

The survival rate was moderate to high (84% to 92%), not differing significantly among the treatments (log rank test, *p* > 0.8), although the antibiotic-exposed animals had slightly higher survival compared to the controls (Figure S1). According to the individual growth curve analysis, the animals exposed to the lowest ciprofloxacin concentration (0.01 mgL^−1^) had a significantly greater maximal body length (BL _max_) compared to the control animals, whereas the K values were similar across the treatments (Figure 1).

**Figure 1.**
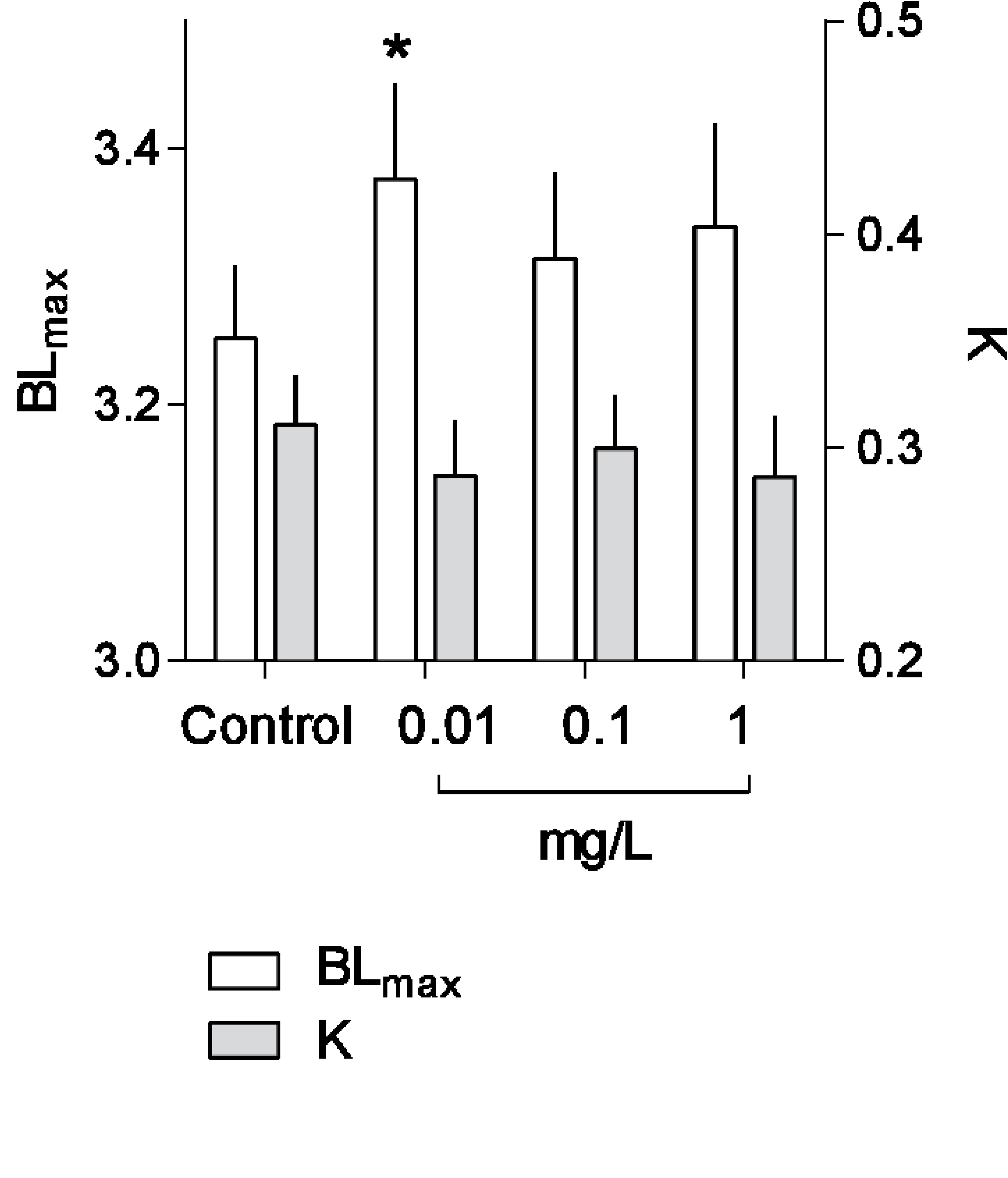
Individual growth curve analysis. Estimated BL_max_ and K values and corresponding 95%-confidence limits for *Daphnia magna* grown in 0.01, 0.1 and 1 mg / L ciprofloxacin and the control.

### Reproduction

The average brood size was significantly higher in all ciprofloxacin treatments compared to the control (GLM, t_263, 267_ = 12.97,*p* < 0.001; Figure 2), with the increase varying from 36% in the 0.01 mg/L treatment (t_263, 267_ = 4.347; *p* < 0.001) to 42% in the 0.1 mg/L treatment (t_263, 267_= 4.05; *p* < 0.001). Also, there was a significant negative effect of time (t_263, 267_ = -2.74; *p* < 0.05), which was mainly related to the low fecundity in the last brood (Tukey HSD, z _(4-1)_:- 3.084, *p* _(4-1)_ < 0.01; z _(4-2)_: -5.97, *p* _(4-2)_ < 0.01; z _(4-3):_ -3.34, *p* _(4-3)_ <0.005). The total number of offspring produced during the experiment per individual female was 27-36% higher in the daphnids exposed to ciprofloxacin compared to controls.

**Figure 2.**
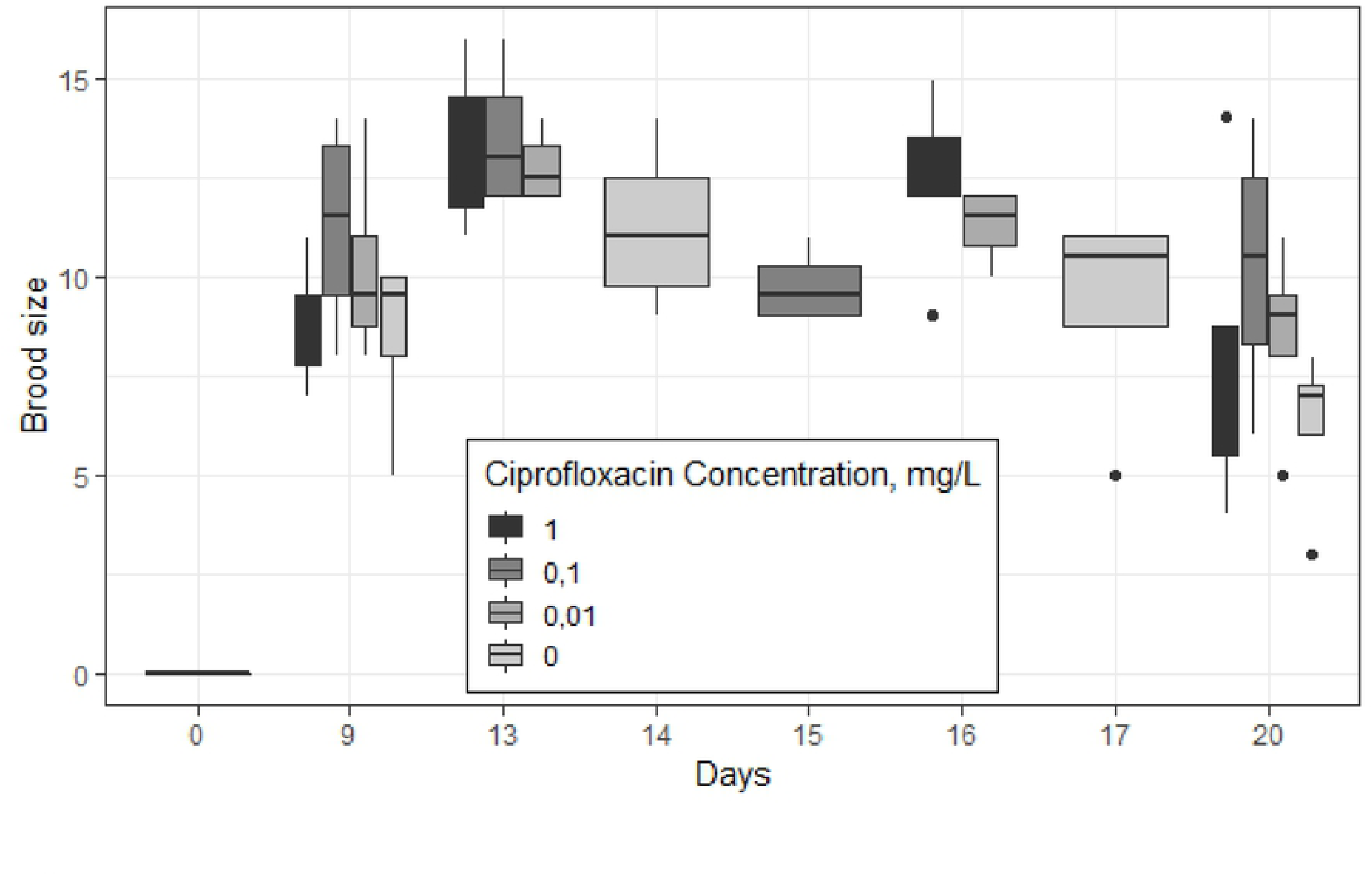
Reproduction of *Daphnia magna* (brood size and time of reproduction) during a 21-d exposure to ciprofloxacin (0.01, 0.1, and 1 mg / L) and a control. Breadth of the box indicates an extended period for clutch release within a treatment, i.e., non-synchronous reproduction. Note that the last clutch was estimated from both the number of offspring released and the number of embryos in the brood chamber at the termination of the experiment.

### Population growth rate

The population growth rate (PGR) varied from 0.26 to 0.30 among the treatments and was higher in the exposed daphnids relative to the control by 17%, 19% and 15 % in the animals exposed to 0.01, 0.1 and 1 mgL^−1^, respectively. The differences from the control were significant for all treatments (Table S1).

### Characterization of the gut microbiota in Daphnia

A total of 1314 high-quality sequences were obtained after trimming and assembly. The core gut microbiome of our test animals was dominated by Proteobacteria, which contributed on average 74% (ranging from 25% to 95% in individual specimens). When all treatments were considered, Actinobacteria (15%), Bacteroidetes (7%), Firmicutes (1%) and Verrucomicrobia (1%) were also common. In the non-exposed animals, the contributions were different, with Proteobacteria, Bacteroidetes and Verrucomicrobia being the most common (Figure S2e). Together, these five phyla formed the core microbiome of the gut and comprised on average 99% of the OTUs assigned to phylum level (Table S5a).

The major classes of bacteria found in all treatments, in order of prevalence, were Betaproteobacteria (35% of total OTUs), Gammaproteobacteria (29%), Actinobacteria (14%), Alphaproteobacteria (9%), Cytophagia (5%), and Verrucomicrobia (1%). In the non-exposed animals, Cytophagia was the third most abundant group, contributing 8 to 36% throughout the experiment, whereas Actinobacteria contributed less than 2% on average. Bacilli, Sphingobacteria and Bacteroidia were found together in about 3% of total reads assigned at class level (Table S5b).

We found members of 62 orders in all treatments (Table S5c). Predominant orders included Burkholderiales (34%), Oceanospirillales (15%), Alteromonadales (10%), Rhizobiales (7%), Micrococcales (5%), and Cytophagales (5%), which was the second most represented order (16%) in the non-exposed animals. The core gut microbiome were formed by these orders along with Propionibacteriales, Corynebacteriales, Pseudomonadales and Methylophilales representing almost 89% of the OTUs assigned at the order level.

Members of 101 families comprising 252 genera were identified as unique reads and assigned at the family and genus level. Across the treatments, Comamonadaceae (33%), Halomonadaceae (15%), Shewanellaceae (10%), and Cytophagaceae (5%) were the most common (Table S5e). In the non-exposed animals, Comamonadaceae (65%) and Cytophagaceae (17%) were the most common. When all treatments were considered, the most abundant genera were *Limnohabitans, Shewanella, Halomonas, Bosea*, and *Leadbetterella*. These genera contributed on average 71% (ranging from 57% to 81%) to the gut microbiota. In the non-exposed animals, however, *Bosea* was not contributing to the core microbiome (Figure S2a).

### The effects of ciprofloxacin on the gut microbiota

Chao1, ACE and Fisher’s alpha indices were negatively co-related to ciprofloxacin concentration (Figure 3a and Table S3 and S4). According to the PCoA, populations exposed to 0.1 and 1 mgL^−1^ clustered closely to each other, with higher loadings on the second axis, which separated them from the control (Figure 4). The homogeneous dispersion (Betadisper, *p* >0.05, Table S3a) met the assumption for further pairwise comparison between the treatments, and a permutation test detected significant differences between the communities exposed to ciprofloxacin and those in control (Permanova, pairwise comparison *p* < 0.05, Table S4). Differential abundance analysis showed the most Ciprofloxacin sensitive bacteria to be *Leadbetterella* (Bacteroidetes), and *Hydrogenophaga* and *Methylotenera*, both Betaproteobacteria. On the opposite end of the scale (most refractory) were *Pseudorhodoferax, Shewanella*, and *Halomonas* (Beta- and Gamma-Proteobacteria), as their abundance in the exposed animals had increased significantly following antibiotic exposure (Figure 5a, Table S6).

**Figure 3.**
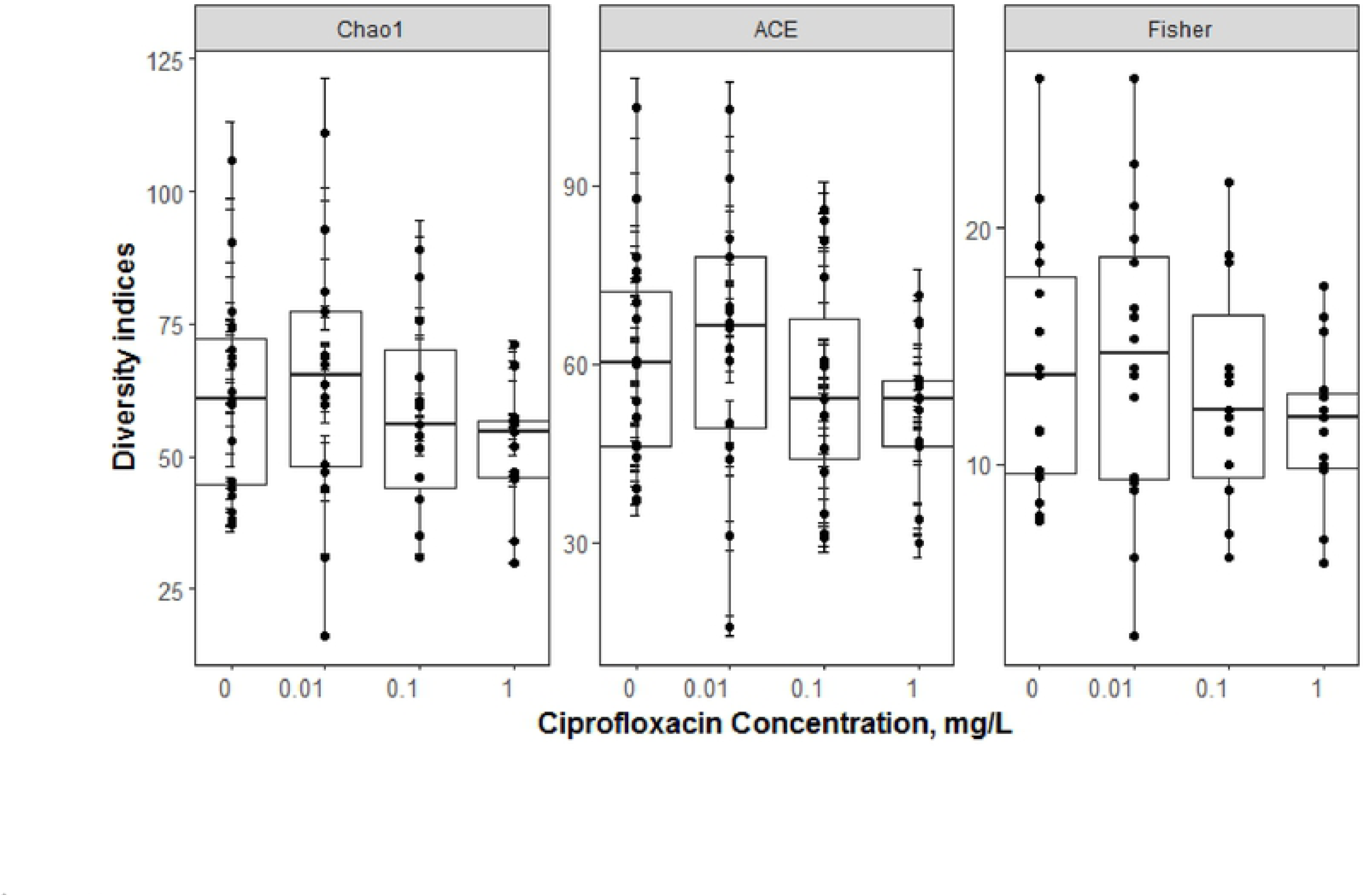
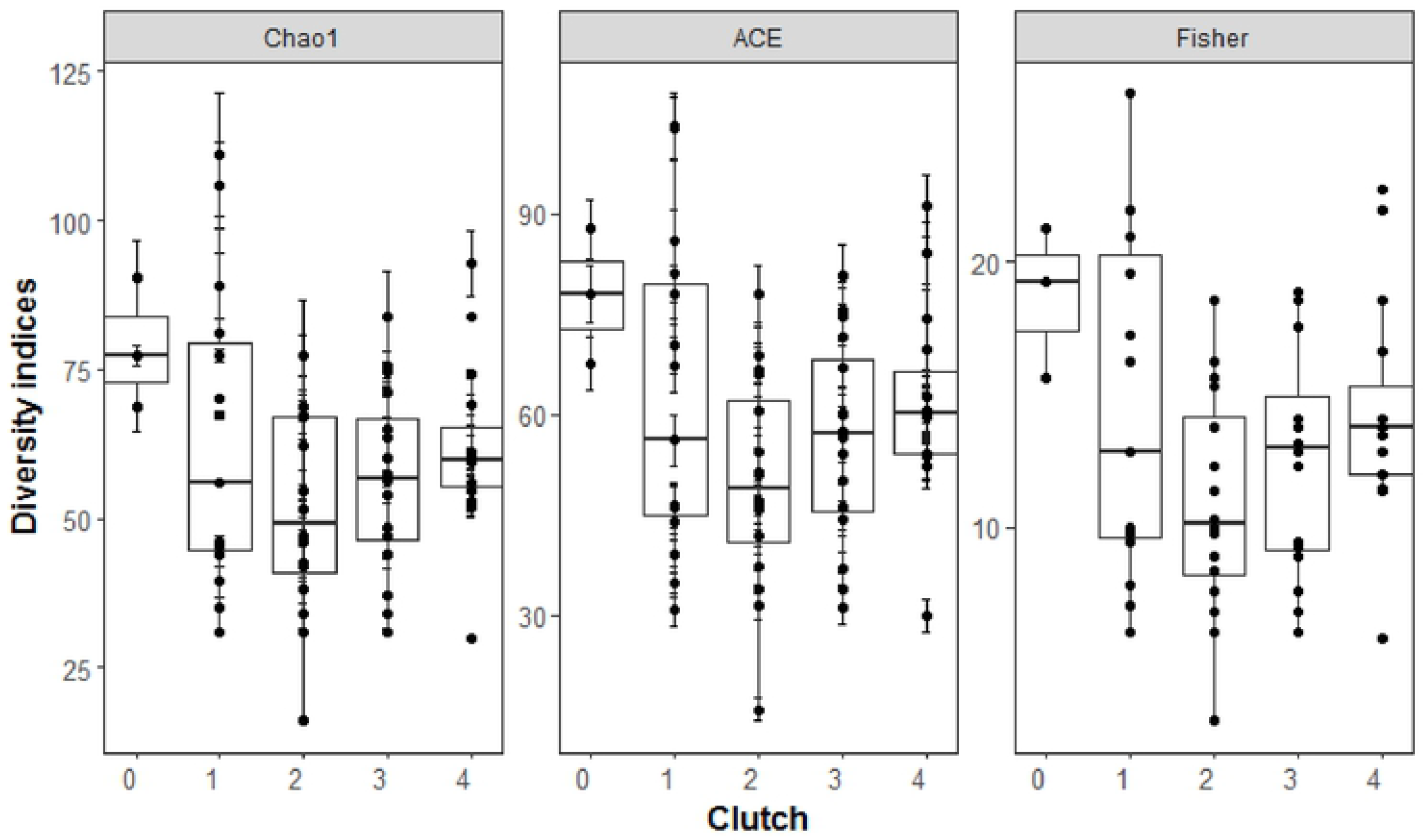
Alpha diversity indices (Chao1, ACE, and Fisher) obtained for gut microbiota. Communities grouped by (a) ciprofloxacin concentration and (b) clutch number during the 21-day exposure. Clutch “0” indicates initial diversity of individuals. Points indicate specific values for individual daphnids.

**Figure 4.**
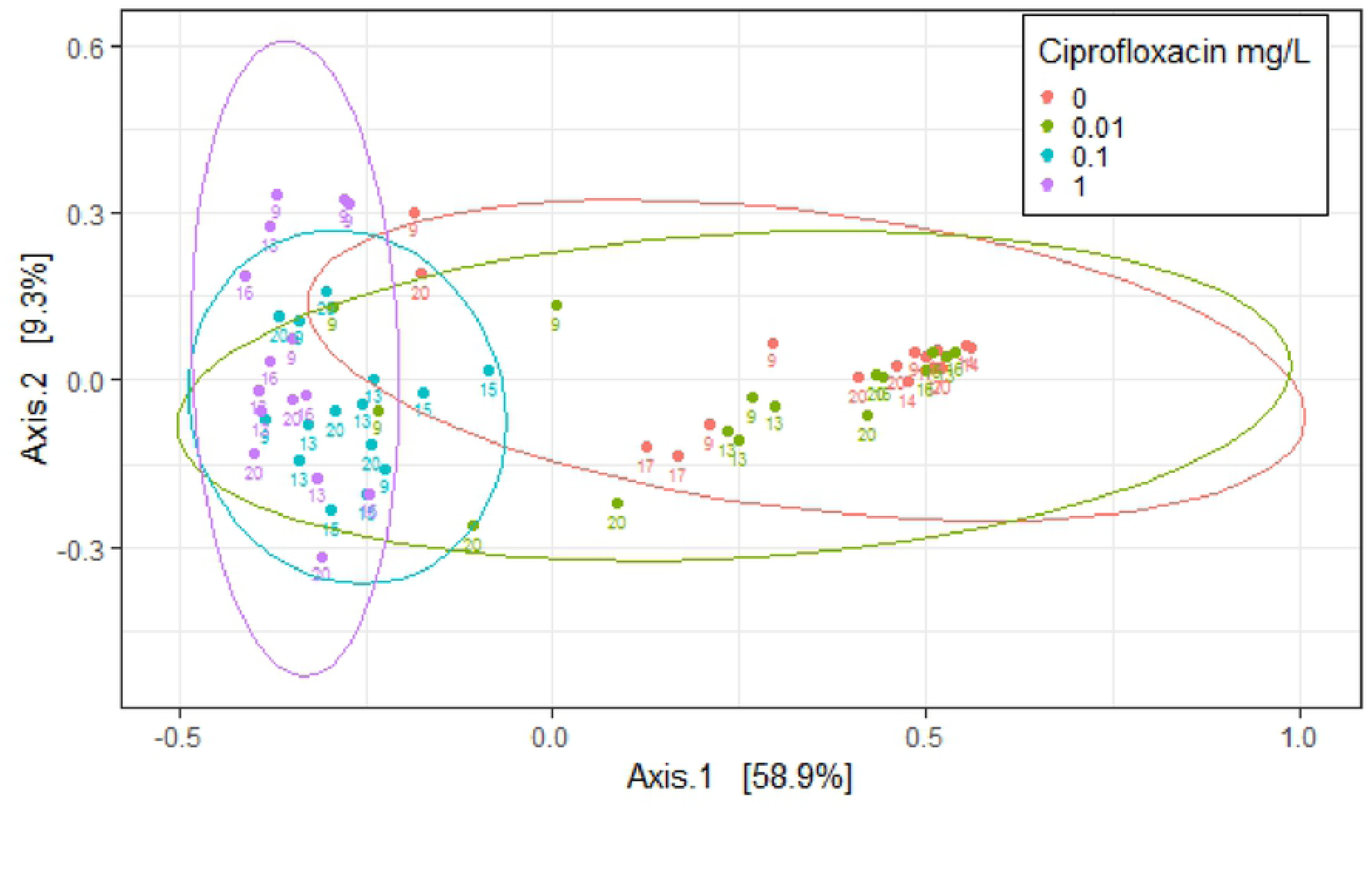
Principal coordinate ordination (PCoA) of the 16S rRNA gene libraries based on the Bray-Curtis dissimilarity. Colors indicate treatments, i.e., concentration of ciprofloxacin (Control: 0, 0.01, 0.1, and 1 mg / L). The ellipsoids represent a 95% confidence interval of normal distribution surrounding each group. Point labels indicate day of sampling. Plot shows the clear clustering of bacterial communities in the treatments exposed to the two highest concentrations of ciprofloxacin (0.1 and 1 mg / L), as well as between communities in the controls and the lowest exposure concentration (0.01 mg / L).

**Figure 5.**
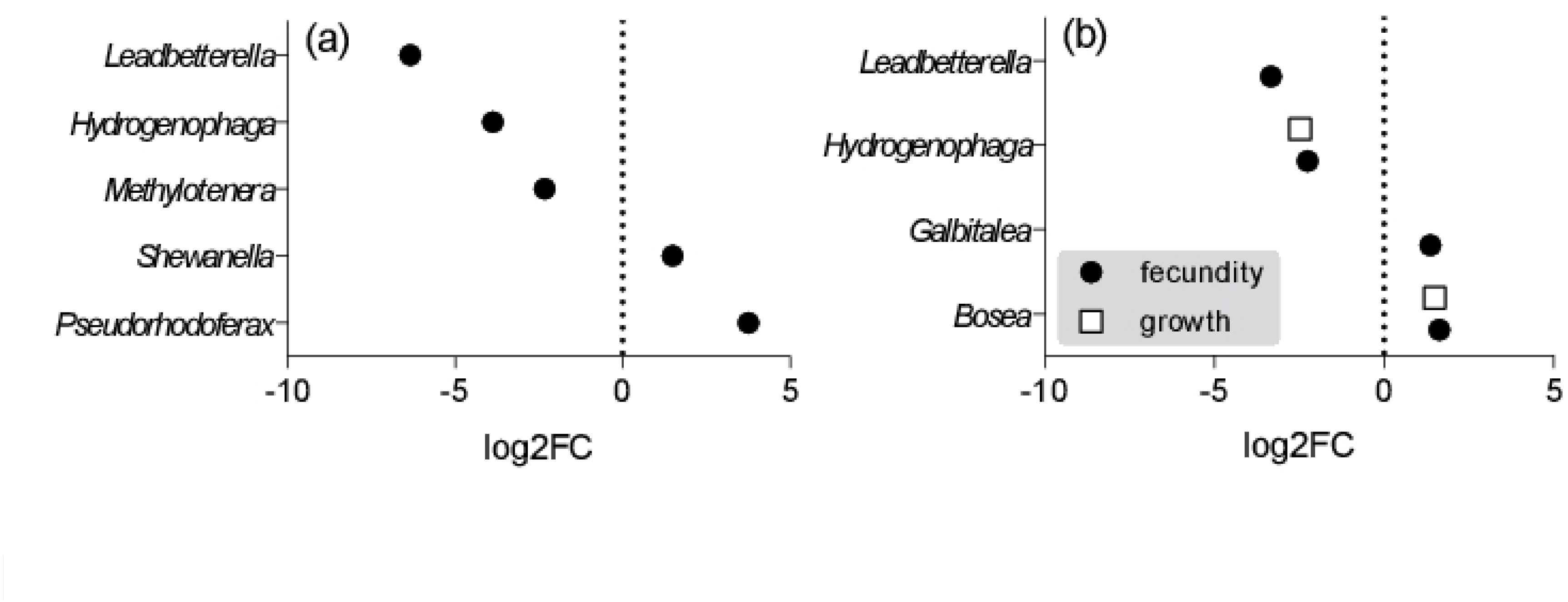
Differential abundance analysis of gut bacteria. Bacterial genera significantly associated with (a) exposure to ciprofloxacin; (b) high somatic growth and fecundity of the host observed during the experiment. The fold change (log2FC) and the associated statistics were derived by the edgeR test.)

### Changes of the gut microbiota with time

Although diversity (Fisher’s alpha) increased with time of exposure, concentration had a more profound than time on this index (Figure 3b; Table S2). Chronic exposure to ciprofloxacin, resulted in a significantly lower diversity in the exposed animals (Figure 3a, Table S2). All diversity indices showed a similar trend over time, with a high diversity during the first two weeks (the first clutch), a decrease observed at the time of the second clutch, following by an increasing trend. However, the time effect was not significant (Table S2).

### Linkages between the gut microbiome and life-history traits

The diversity indices correlated negatively with fecundity, while only Fisher’s alpha had a positive correlation with body size. The differential abundance analysis indicated that genera *Bosea* and *Hydrogenophaga* were more abundant in the daphnids with high and low somatic growth, respectively (Table S7; Figure 5b). Moreover, *Bosea* and *Galbitalea* were significantly more abundant in the daphnids with higher fecundity, whereas abundances of *Leadbetterella* and *Hydrogenophaga* in these individuals were significantly lower (Table S7, Figure 5b). Thus, *Bosea* and *Hydrogenophaga* were consistently associated with high and low growth phenotypes, respectively.

### Biomarker ORAC/Protein responses to antibiotic exposure

The total antioxidant capacity (ORAC, g Trolox eq. g protein^−1^) was significantly higher in the animals exposed to lower concentrations of ciprofloxacin (0.01 and 0.1 mgL^−1^) (Figure 6, Table S8). Moreover, there was a significant positive relationship between the individual ORAC and body length (GLM; Wald stat. = 5.83, p < 0.02; Table S9, Supporting Information) across the concentrations tested. The correlations between the ORAC values and diversity indices were negative and marginally significant (Table S10, Supporting information).

**Figure 6.**
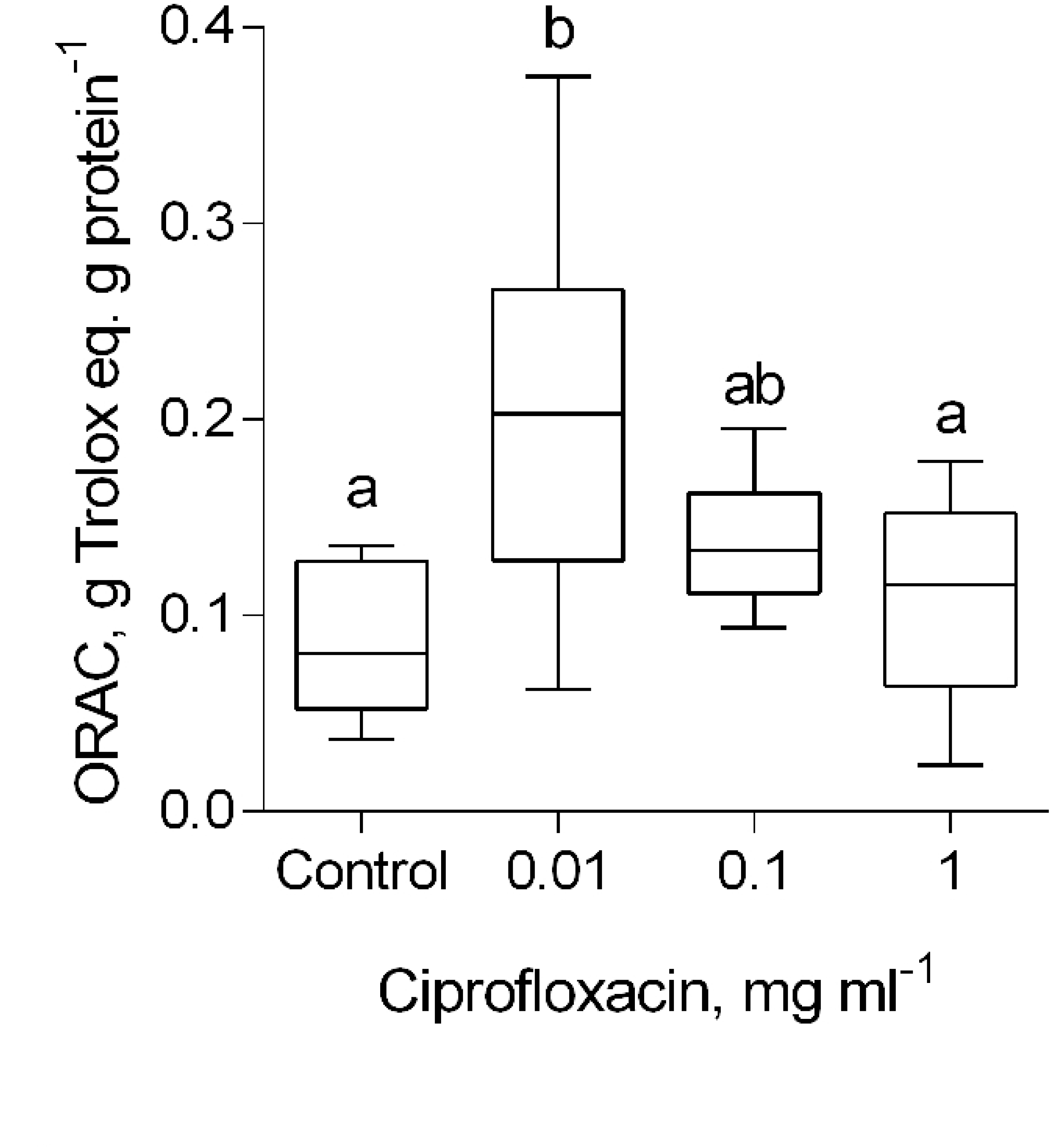
*Daphnia magna*: response of the total antioxidant capacity (ORAC, g Trolox eq. g protein^−1^) to the ciprofloxacin concentration. The individuals sampled after their fourth clutch were excluded, because some of them contained eggs in the brood chamber. The non-matching letters indicate significant differences between the groups (Tukey’s multiple comparisons test; p < 0.05). See Table S9 for details on the statistical comparisons.

## Discussion

The intestinal microbiome plays an essential role in regulating many aspects of host physiology, and its disruption through antibiotic exposure has been implicated in microbiota-mediated consequences on host fitness. We examined effects of chronic antibiotics exposure on *Daphnia magna* gut microbiota in concert with fitness-related responses of the host. As hypothesized, the exposure to ciprofloxacin resulted in profound changes in the microbiome and a reduced microbial diversity at all concentrations tested (0.01 to 1 mgL^−1^). Surprisingly, no negative effects on daphnid antioxidant levels, fitness and mortality were observed. Moreover, the negative changes in the microbiome coincided with increased antioxidant capacity, individual growth and host reproduction and, as a result, significantly higher population growth in the animals exposed to ciprofloxacin. Thus, the hypothesized positive correlation between microbiome diversity and host performance was not observed. Our findings indicate that reliance on shifts in taxonomic composition of bacterial community generates an incomplete picture of the functional effect of antibiotic intervention in a non-target eukaryote. A full mechanistic understanding will require further study of the specific functional relationships between the host and its core microbiome, and the integration of metabolomic and phenotypic data. Moreover, in case of antibiotic-mediated intervention, we need to disentangle direct effects of the exposure on host physiology. This is already evident in human microbiome study where drug effects on mitochondrial activity are known to confound (38,39).

### *Core microbiome of* Daphnia magna

Proteobacteria, Actinobacteria and Bacteroidetes comprise a core microbiome of the *Daphnia magna* intestine. Most taxa (or their close relatives) identified in this study as a part of core microbiome have previously been reported in *Daphnia* (21,40,41). The Comamonadaceae family of Burkholderiales have been shown to be the most abundant family in *Daphnia* gut microbiota (41,42), and were most prevalent in our test animals. Other taxa found in high abundance were the Gammaproteobacteria orders Oceanospirillales and Alteromonadales, and the families *Nocardioidaceae, Microbacteriaceae*, and *Moraxellaceae* (21,12). On the genus level, more differences between earlier reported daphnid associated taxa and our dataset were evident. In addition to *Limnohabitans*, other identified microbial taxa were *Pseudorhodoferax* and *Hydrogenophaga* (Burkholderiales) but <u>not</u> the previously reported *Bordetella, Cupriavidus* (43), *Ideonella* and *Leptothrix* spp. (41). Also, *Enhydrobacter* was the dominant genus of Moraxellaceae in our study (Table S5e), while *Acinetobacter* spp. was reported in other studies (12,20). *Methylibium* was only found in the animals that were exposed to 0.01 mg / L of Ciprofloxacin and not in the non-exposed individuals, suggesting that this genus is relatively rare if normally present. Together, our results present a relatively stable bacterial composition in the *Daphnia* gut from a higher taxonomic level, suggestive of functional or other redundancy in the preferred association of daphnids with their microbiota components.

### Effects of Ciprofloxacin on the Daphnia gut microbiome

Drug exposure significantly altered the microbiome, with a decrease or even the disappearance of many taxa by the end of the experiment at lowest exposure concentration and within a first week at higher concentrations (Figure 3b, Table S5). Although diversity decreased with both ciprofloxacin concentration and exposure time, only the concentration effect was significant (Table S2, Figure 3). G+ bacteria, mostly *Actinobacteria* and *Firmicutes*, were better able to withstand ciprofloxacin effects as their relative abundance increased with drug concentration (Figure. S4a), while the G-bacteria had divergent responses (Figure. S4b). For example, *Hydrogenophaga* and *Pseudorhodoferax*, both belonging to the G-genus *Burkholderiales*, had clearly opposite responses, decreasing and increasing, respectively, with increasing concentration. This is in line with earlier studies that demonstrated higher susceptibility to Ciprofloxacin among G-bacteria, as compared with co-occurring G+ species (44). This is evident for the typically low minimum inhibitory concentrations, MICs, estimated for Alphaproteobacteria, such as Escherichia/Shigella, (commonly in the low µM range) as compared with that for many Firmicutes, which are usually in the mM range.

At higher concentrations of Ciprofloxacin, several genera representative of the core microbiome declined to non-detectable levels; the *Limnohabitans* genus was replaced by *Halomonas* and *Shewanella*, whose relative abundances increased with drug concentration (Table S5e). *Shewanella* is a known acid producer (45) and may alter the pH balance in the gut microenvironment when at higher densities. This would suppress the growth of *Limnohabitans* who grow preferentially under neutral and alkaline conditions (46). Such community-level effects probably play a significant role in the dynamics of specific bacterial taxa as a result of the exposure to antibiotics.

### Effects of Ciprofloxacin on Daphnia life history traits and antioxidant levels

Previous studies on aposymbiotic daphnids showed that disruption in gut microbiota, either by drugs or diet, had adverse effects on nutrition (40) (11), immunity (8), growth (12), fecundity (22), and longevity (47). The effects that we observed however, were most prominent at low antibiotic concentrations. Despite the ciprofloxacin-induced shifts in the microbiome composition, ORAC levels, growth and reproduction in the daphnids were similar or even significantly higher than in controls. The discrepancy between the microbiome and the organism-level responses may result from the variable susceptibility of various microbes to the broad-spectrum Ciprofloxacin and additional variability related to induction of the SOS response pathways in different taxa.

The mismatch between microbiome change and host response suggests that other drivers, such as a direct effect of Ciprofloxacin on the host, were involved, leading to the observed effects on growth and reproduction. In line with this, a biphasic dose-response to ciprofloxacin observed in human fibroblast cells, manifesting as increased cell proliferation and viability when compared to non-exposed controls (48). In *Daphnia magna*, the reproduction response to ciprofloxacin was also biphasic, with stimulatory effects at concentrations below 5 mg/L (49). This is in line with the positive response induced by the test concentrations utilized in our study (0.01-1 mg/L). In mice, ciprofloxacin has also been shown to improve survival by enhancing immune efficiency via stimulating cytokine production (50). In addition, several *in vitro* and *in vivo* studies using animal and tissue models have revealed that fluoroquinolones like ciprofloxacin, induce oxidative stress via reactive oxygen species (ROS) production, in a dose- and time-dependent manner (49,51).

Measurable ROS production was observed following an exposure to ciprofloxacin at concentrations as low as 0.025 mM (53), which is within the concentration range used in our study. At low levels of such pro-oxidative exposure, the additional production and/or activity of the endogenous antioxidant enzymes and low-molecular weight antioxidants to remove the continuously generated free radicals would increase (54). In the daphnids exposed to the lowest Ciprofloxacin concentration, a significant increase in ORAC levels (Figure S3) suggests that exposure had direct stimulatory effects on the antioxidant production. Moreover, we observed a positive correlation between the ORAC levels and animal body size across the treatments indicating a possible primary mechanism behind the observed effects being a hormetic shifting of redox environment by pro-oxidative ciprofloxacin, antioxidant response and the resulting beneficial effects on growth. Such effects are in agreement with a concept of physiological conditional hormesis (55) and suggest a possible mechanism for the direct response of *Daphnia magna* to Ciprofloxacin exposure at environmentally relevant concentrations. An important caveat is that hormesis, also shown to occur in several microbes’ response to quinolones and fluoroquinolones (the so-called paradoxical effect) (56) might be universal and thus ciprofloxacin may be suboptimal for the uncomplicated study on microbiome involvement following dose-response relationships.

### Microbiome-fitness relationships

Although elevated growth and reproduction were associated with some bacterial taxa, there was no clear signal for involvement of the gut microbiome in the high-growth phenotype. This is suggestive of a form of redundancy in host-microbiome function, i.e., microbes can be exchanged with little or no penalty. Moreover, as mechanisms governing most observed associations are not well understood, definitive conclusion of direct effects by specific microbes is intuitively discouraged. In particular, several taxa (*Bosea* and *Shewanella*) significantly associated with fitness-related variables have been shown to be highly resistant to ciprofloxacin (57,58). Thus selection although acting directly on the polymicrobial community, it does so differentially and although the effect may be due to absolute numbers of microbes, the cumulative physiological and metabolic state may matter more. In line with this, the relative abundance of those genera that were associated with higher fecundity and growth barely comprise 5% of the organism’s core microbiome (Table S5), suggesting that sheer abundance was unlikely to be the primary factor driving host fitness.

An important caveat is that hormesis, also shown to occur in several microbes’ response to quinolones and fluoroquinolones (the so-called paradoxical effect) (56) might be universal and thus ciprofloxacin may be suboptimal for the uncomplicated study on microbiome involvement following dose-response relationships.

It is a common view that strains capable of supplying essential elements for reproduction and growth would benefit the host. For example, the key components of *Daphnia* gut microbiota, *Limnohabitans, Aeromonas* and methanotrophic bacteria (47), have been linked to acquisition of essential amino acids (58,38), polyunsaturated fatty acids (PUFA) and sterols (60) that positively affect *Daphnia* growth and reproduction (9,61). Surprisingly, none of these taxa were associated with elevated growth and fecundity in our study. This also speaks for functional redundancy although additional studies would be required to show this. At the genus level, only *Bosea* and *Galbitalea* had significantly positively association with *Daphnia* growth and fecundity, whereas *Leadbetterella* and *Hydrogenophaga* correlated negatively. *Leadbetterella* and *Hydrogenophaga* were previously found to be associated to 8 Daphnia genotypes (62). More interestingly however, the Bradyrhizobiaceae (*Bosea*) and Microbacteriaceae (*Galbitalea*) are bio-degraders capable of producing hydrolytic enzymes such as chitinase, cellulase, glucanase, protease, etc. (57,63). As these are positively correlated with fecundity and host fitness, it suggests that increased network density and number of degradation pathways may contribute by providing essential nutrients from more available substrates (64). Regardless of the mechanisms underlying their increased abundance, resistance, or at the very least, refractoriness to Ciprofloxacin cannot be ignored. Such effects would be evident in perturbed outcome of inter- and intra-species competition and illustrates one of the difficulties facing future studies into host-microbiome interactions.

## Acknowledgements

The computations were performed on resources provided by SNIC through Uppsala Multidisciplinary Center for Advanced Computational Science (UPPMAX) under project 2018/8-68. Sequencing and analysis of microbiome results were made possible by grant # 20160933 from the Stockholm County Council (SLL) to KU.

## Supporting information

**Figure S1**. Survival of *Daphnia magna* exposed to ciprofloxacin (0.01, 0.1 and 1 mg/L) and in the control (M7 only) during the 21-d exposure.

**Figure S2**. Relative abundance of bacterial taxa in the microbiome of *Daphnia magna* from the non-exposed treatments: (a) genera, (b) families, (c) orders, (d) classes, and (e) phyla. The data are grouped by the exposure week, 1 to 4 (Y-axis). Animals collected at the termination of the experiment are included in the week 4.

**Figure S3**. Change in the total antioxidant capacity (ORAC, g Trolox eq. / g protein) in individual daphnids during the course of the experiment. The data are shown for each treatment (ciprofloxacin exposure, 0.01, 0.1 and 1 mg / mL) and the control. The regression line and the 95%-confidence interval are shown to indicate the overall direction of change over time; no trends are significant (p > 0.05).

**Table S1**. Population growth rate (*r*) of *Daphnia magna* in the control and ciprofloxacin exposure (0.01 – 1 mg/L) and the corresponding 95-% confidence interval estimated by bootstrapping. Asterisk indicates significant difference from the control; when the confidence intervals were not overlapping, the difference was considered significant.

**Table S2**. Diversity indices were calculated using individual data rarefied to equal sequencing depth at treatment level. Effects of concentration and time on the diversity indices (Fisher’s alpha, Chao1 and ACE) were evaluated using GLM. Interactions were included first in each model but omitted when found not significant.

**Table S3**. Multivariate homogeneity of groups’ dispersions (betadisper) of samples analyzed according to treatment (Ciprofloxacin concentration).

**Table S4**. PERMANOVA output with Bray-Curtis dissimilarity testing differences between treatments at family level.

**Table S5**. Relative contributions of the ten most common bacterial taxa to gut microbiota of *Daphnia magna* exposed to ciprofloxacin (0.01. 0.1. and 1 mg/L) and in control (0 mg/L) as well as the average relative abundance for all treatments.

**5Table S6**. Differential abundance of individual genera estimated by edgeR-function and testing taxa-specific responses to ciprofloxacin exposure. The positive log2FC values indicate increased relative abundance in the exposed daphnids compared to the controls. Significance presented at false discovery rate of 5%. (FDR<0.05). See also Figure 7a.

**Table S7**. Differential abundance analysis of individual genera estimated by the edgeR-function and testing associations between the microbiome and host fitness. The genera positively associated with high growth or fecundity of *D. magna* have positive log2FC values. All values reported are significant at false discovery rate of 1%. (FDR<0.01). See also **Error! Reference source not found.**b.

**Table S8**. Effect of ciprofloxacin concentration (mg mL^−1^) on antioxidant capacity in *Daphnia magna*: (A) ANOVA results testing overall effect, and (B) Pair-wise comparisons using Tukey’s multiple comparisons test; p < 0.01: **, p < 0.05: *; and p > 0.05: ns. The individuals sampled at the termination of the experiment were excluded, because some daphnids contained eggs in the brood chamber. As the reference group, we used the daphnids exposed to the highest concentration. See also Figure S3.

**Table S9**. Generalized linear model output linking antioxidant capacity to daphnid body length across the concentrations tested. Normal error structure and log-link function were applied. The animals collected at the termination of the experiment were excluded, because they had eggs in the brood chamber, which may affect the ORAC values.

**Table S10**. Spearman rank correlation between the ORAC values in the daphnids and diversity indices of their gut microbiome.

**Appendix S11.**

R script used to calculate population growth rate of daphnids applying Euler-Lotka equation:

